# Transcriptomic analysis of *Lentinus squarrosulus* provide insights into its biodegradation ability

**DOI:** 10.1101/2020.09.28.316471

**Authors:** Aarthi Ravichandran, Atul P Kolte, Arindham Dhali, S. Maheswarappa Gopinath, Manpal Sridhar

**Affiliations:** ICAR-National Institute of Animal Nutrition and Physiology, Adugodi, Bengaluru 560 030, India; Women Scientist, Department of Science & Technology (Govt. of India); Department of Biotechnology, Davanagere University, Davanagere, India

**Keywords:** *Lentinus squarrosulus*, biodegradation, transcriptome, lignocellulose

## Abstract

Basidiomycetes are of special interest in biotechnological research for their versatile potential in degradation of lignocellulosic biomass. This study accordingly reports analysis of transcriptome of a white-rot Basidiomycete *L.squarrosulus* grown in simple potato dextrose broth supplemented with aromatic compound, reactive black dye to gain an insight into the degradation ability of the fungus. RNA was sequenced using Illumina NextSeq 500 to obtain 6,679,162 high quality paired end reads that were assembled *de novo* using CLC assembly cell to generate 25,244 contigs.Putative functions were assigned for the 10,494 transcripts based on sequence similarities through BLAST2GO 5.2 and Function annotator. Functional assignments revealed enhanced oxidoreductase activity through the expression of diverse biomass degrading enzymes and their corresponding co-regulators. CAZyme analysis through dbCAN and CUPP revealed the presence of 6 families of polysaccharide lyases, 51 families of glycoside hydrolases, 23 families of glycoside transferases, 7 families of carbohydrate esterases and 10 families of Auxiliary activities.Genes encoding the ligninolytic enzymes and auxiliary activities among the transcript sequences were identified through gene prediction by AUGUSTUS and FGENESH. Biochemical analysis of a couple of biomass degrading enzymes substantiated the functional predictions. In essence, *L.squarrosulus* grown in a simple medium devoid of lignocellulosic substrate demonstrated presence of a repertoire of lignocellulose degrading enzymesimplying that source of lignocellulose is not required for expression of these biomass degrading enzymes. The study hereby underlines the significance of *L.squarrosulus* in biomass degradation and its future functional exploitation in biomass conversion applications.

## Introduction

Basidiomycetes have been the focus of intense research by mycologists for their antioxidant, antiparasitic, immune modulating effects and considerably on biomass degradation [1]. Biodegradation of plant biomass especially cellulose and hemicellulose have been examined with different groups of Basidiomyceteshowever,the distinctive ability of depolymerization of lignin is an attribute of relatively few species.This group of basidiomycetous fungi commonly referred to as “white-rot” is saprophytic on plant organic matter and they acquire energy from plant polysaccharides cellulose and hemicellulose by disrupting the lignin complex, a refractory polymer cementing the carbohydrates [2].The biological and economic significance of lignin degradation is exemplified by the multifarious downstream applications of the plant polysaccharides. White-rot fungi secrete ligninolytic enzymes for degrading lignin that includes phenol oxidases like laccases and peroxidases like lignin peroxidase, manganese peroxidase and versatile peroxidase[3]. Ligninolytic enzymes bank upon H_2_O_2_ producing enzymes and other intermediate products of lignin degradation which inturn serve as a cofactor for further depolymerization. Enzymes entailed in decomposition of lignocelluloses by these fungi is not limited to ligninolytic enzymes but includes exo, endo β-glucanases, xylanases, esterases, pectinases, mannanases, cellobiohydrolases, polysaccharide monooxygenases to mention a few.

Production and expression of these enzymes is regulated chiefly by the environmental conditions and substrate of growth [4].Having stated the complexity of lignocellulose degradation, relatively little is known about this metabolic process and the subsequent polysaccharide degradation. Hence in this arena, transcriptomic analysis of the lignicolous fungi grown on different substrates was attempted by researchers worldwide to gain more insight into the degradation process [5–9]. Nevertheless, these studies have so far focused on fairly small group of specieswherein the expression of fungal genes in response to lignocellulosic biomass as substrate was reported. Our work on these lines was aimed to delineate the expression of lignocellulose degrading genes of *L. squarrosulus* grown in a simple medium devoid of lignocellulosic substrate. Lignocellulosic biomass is complex network of polysaccharides cemented with lignin and hence induces expression of plethora of biomass degrading enzymes in the colonized white-rot. To better understand the regulation of these enzymes on a non-lignocellulosic substrate, attempts were madeto investigate the transcriptome of *L.squarrosulus* cultured in potato dextrose broth induced with reactive black 5 for ligninolytic enzymes. *L.squarrosulus* is a tropical white-rot fungus that grows immensely on decaying wood and saw dust. *L.squarrosulus* is a rich producer of ligninolytic enzymes predominantly versatile peroxidase as identified through our study.Though complete depiction of the ligninolytic machinery is very far, transcriptomic analyses of this fungus grown in synthetic medium can add significant insights to the ligninolytic process.

## Materials and methods

### Fungal strain and culture conditions

The fungus subjected in the study is a wild isolate from Western Ghats, India determined through ITS sequencing. This isolate identified as *L.squarrosulus* was propagated through tissue culture and maintained by frequent sub-culturing on potato dextrose agar at 28°C. Production of ligninolytic enzymes were induced by supplementing potato dextrose broth with 0.01% reactive black 5 (RB5), an azo dye.Erlenmeyer flask with 50 ml of culture medium was inoculated with homogenized mycelia at 2% from freshly grown seed culture of the fungus. The flasks were continuously agitated at 100 rpm and incubated at 28±2 °C for seven days.

### Enzyme activitymeasurements

*L.squarrosulus* cultivated in potato dextrose broth induced with RB5 for production of ligninolytic enzymes was assessed for existence of extracellular biomass degrading enzymes.Extracellular biomass degrading enzymes like cellulase, xylanase, polygalacturonase, mannanase, α-glucosidase, β-glucosidase and xylan esterases were assayed in the culture supernatant besides ligninolytic enzymes of laccase, manganese peroxidase and versatile peroxidase.Culture supernatant was filtered to remove mycelia and utilized for biochemical assays.

Cellulase activity was deduced with carboxymethyl cellulase as substrate. The assay mixture consisted of substrate 0.25% and 0.2 ml culture supernatant in 50mM phosphate buffer pH 6.8 [10]. The procedure for xylanase estimation is similar to cellulase with the substrate being 0.06% oat spelt xylan. Polygalacturonase assay reaction mixture comprised polygalacturonic acid 0.45%, culture supernatant 0.1 ml in 50 mM acetate buffer pH 5 [11]. Mannanase activity was inferred with 0.5% locust bean gum as substrate. 0.1 ml of culture supernatant was reacted with 0.9 ml substrate solution in phosphate buffer pH 6.8 [12].Reducing sugars produced through theenzyme reaction was measured by dinitro salicylic acid (DNS) method at 575 nm [13]. D-glucose, D-xylose, D-galacturonic acid and D-mannose were used as standards for estimation of cellulase, xylanase, polygalacturonase and mannanase activities respectively.

α-glucosidase, β-glucosidase and xylan esterases activities were interpreted using 0.1% p-nitrophenol α-D glucopyranoside, 0.1% p-nitrophenol β-D glucopyranoside and 0.02% p-nitrophenol acetate as substrates correspondingly with p-nitrophenol as standard [10,14]. The substrates were accordingly dissolved in 0.1M phosphate buffer pH 6.8 prior to use and reacted with 0.1 ml of culture supernatant as crude enzyme sample. Enzyme activity was determined by quantifying the amount p-nitrophenol released from the reaction at 400 nm.

1.6 mM ABTS in 100 mM sodium acetate buffer (pH 4.5) was used for estimation of laccase activity (ε_420_ 36000 M^−1^ cm^−1^). Peroxidases interference with laccase activity was corrected using 0.5μg/ml catalase in the reaction mixture. Manganese oxidation activity was deduced using 0.5 mM MnSO_4_ and 100 mM malonate buffer pH 4.5. Formation of Mn^3+^ malonate complex was measured at 270 nm.Decolorization of RB5 was determined in 100 mM sodium tartrate buffer pH 3 with 10 μM of RB5. The reaction was monitored through decrease in absorbance at 598 nm (ε_598_24000 M^−1^ cm^−1^). Reaction for determination of peroxidases were initiated with 0.1mM H_2_O_2_.

### RNA isolation and sequencing

Fungal biomass was filtered from the culture supernatant on day-7 during the late log phase and ground to fine powder using liquid nitrogen. Total RNA was isolated using Genetix RNA sure Plant minikit according to manufacturer’s protocol.Concentration of RNA was measured on nanodrop spectrophotometer and quality assessed bydenaturing agarose gel electrophoresis. 1 μg of total RNA was used for analysis wherein mRNA was enriched from total RNA using poly T attached magnetic beads. Sequencing libraries were prepared using TruSeq stranded library preparation kit and quality assessed through Agilent 4200 Tape station.The library was then subjected to paired end sequencing in Illumina NextSeq 500 platform.

### Reads mapping and annotation

Raw reads were processed in Trimmomatic v0.35 for removing adapter sequences, ambiguous and low quality sequences. The quality reads were then assembled in CLC assembly cell. Annotation was performed using BLAST2GO 5.2 software for identification of Gene Ontology (GO) terms [16]. Exhaustive functional annotations of the transcript sequences were performed through Function annotator [17]. Function Annotator assisted in annotation of enzymes, best hits to NCBI nr database, GO terms, conserved domain, transmembrane protein, lipoprotein peptide, signal peptide and also subcellular localization.KEGG orthology designation was obtained based on homology searches in KAAS [18].

### CAZyme annotation

The catalytic and other functional domains of enzymes involved in metabolism and transport of carbohydrates were elucidated based on signature domains of each CAZy families through dbCAN2 meta server [19]. Short sequence reads were submitted to dbCAN2 meta server for automated CAZyme annotation using tools HMMER, DIAMOND and Hotpep. CAZyme domain HMM database is referred for identification of CAZyme domain boundaries whereas DIAMOND is used for fast homology based searches to the CAZy database. Hotpep identifies conserved motifs in Peptide Pattern Recognition (PPR) library. In addition, Conserved Unique Peptide Patterns in the CAZymes placed them in different functionally related protein groups (CUPP).Finer level of classification based on the similarity of protein sequence and peptide signature of CUPP group was used to classify protein families of the transcript sequences [20].

### mycoCLAPAnalysis

Further functional analysis of lignocellulose active proteins were acquiredthroughmycoCLAP [21]. mycoCLAP database contains comprehensive information on fungal and bacterial genes encoding lignocellulose acting proteins that were characterized through biochemical studies.Transcript sequences generated were subjected to homology search against the mycoCLAP sequences through BLASTfor significant insight into the lignocellulose acting proteins present in the transcriptome.

### Gene prediction

Gene identification was accomplished through AUGUSTUS web server, a gene prediction software based on generalized hidden markov model [22]. The short read nucleotide sequences were fed as input to the AUGUSTUS web server for gene predictions based on *Phanerochaete chrysosporium,* an another White-rot genome. Parameters were set to detect genes from both strands and also through alternate splicing. Another *ab initio* gene prediction software FGENESH based on hidden markov model was also used for gene identification [23]. Sequences were uploaded to FGENESH for gene prediction using organism specific gene finding parameters of *Trametes cinnabarina,a* white-rot that displayed significant nucleotide sequence homology in Function Annotator.

### Data Availability

The raw sequencing data referred in this project was submitted to the NCBI Sequence Read Archive under the accession number PRJNA640439 (Release date: 2021-07-01).

## Results

### Enzyme activity measurements

*L.squarrosulus* was grown in simple synthetic medium with Reactive Black 5 (RB5), an azo dye as inducer for the production of ligninolytic enzymes.Our previous work on ligninolytic enzymes demonstrated that these enzymes were strongly stimulated in presence of aromatic compounds and lignocellulose substrates. In particular, the recalcitrant dye RB5 stimulated higher titers of ligninolytic peroxidases. Lignin degrading enzymes of this fungus particularly versatile peroxidase, manganese peroxidase and laccase were strongly induced in presence of the azo dye. Manganese oxidizing peroxidase activity increased to a maximum of 10 U/ml in induced medium compared to the uninduced medium besides laccase activity. Biomass degrading enzyme activities of cellulase, xylanase, polygalacturonase, mannanase, α-glucosidase, β-glucosidase and xylan esterase were studied to obtain deeper understanding on the regulation of these enzymes comparatively to the ligninolytic enzymes. Only trivial α-glucosidase activity was observed in induced and uninduced media. The interference of the mannanase substrate, locust bean gum with glucose in the culture supernatant impeded the detection of this enzyme.

Rise in cellulase activity was observed on day-7 with activity higher in uninduced than induced medium. The same trend was observed with xylanase, acetyl esterase and polygalacturonase whereas β-glucosidase activity was higher in induced medium. Though the instance of maximum activity and the duration varied for each enzyme, the performance of uninduced medium was optimal for production of biomass degrading enzymes other than ligninolytic enzymes. Ligninolytic enzymes activity was higher in induced medium attributed to complex aromatic nature of the azo dye. This evidently signified that the specific activity of ligninolytic enzymes were higher in induced medium.

### RNA extraction and sequencing

cDNA sequencing libraries were prepared from total RNA of late log phase cultures of *L.squarrosulus* grown in RB5 induced potato dextrose broth.10 million reads of data was generated from initial RNA concentration of 125 ng/μl with mean fragment size of 447 bp. 6,679,162 high quality paired end reads were obtained after removal of adaptor sequences and trimming of low quality bases.The raw reads were aligned *de novo* through CLC assembly cell and 25,244 contigs were obtained.

### Functional annotation

Annotation of the sequences based on similarity was performed using BLAST2GO. BLAST2GO determined the putative functions of the transcripts and categorized them by biological process, cellular component and molecular function. To further explore the function and significance of the transcripts, the sequences were analyzed throughFunction annotator. Function annotator presented efficient interpretation of the reads on GO terms, enzyme groups, domain identification, sub-cellular localization, protein secretion, transmembrane proteins and also on the taxonomic relationship at different levels. Sequence similarity with NCBI non-redundant database yielded equivalent sequences with lowest e-values.

The taxonomic information furnished based on the proportion of similarity sequences placed *L.squarrosulus* close to *Dichomitus squalens* at the species level. The subsequent best hits were *Trametes versicolor* and *Trametes cinnabarina* as depicted in Figure 2. Polyporaceae are major group of white-rot fungi with *L.squarrosulus, Dichomitus squalens* and *Trametes versicolor* belonging to the core polyporoid clade, one of the major clade with rich catabolic ability under Polyporales. Analysis based on ITS, RNA polymerase II and nrLSU sequences support the similarity of *Lentinus* to *Dichomitus* and *Trametes* at the genus level [24].

**Figure 1:**
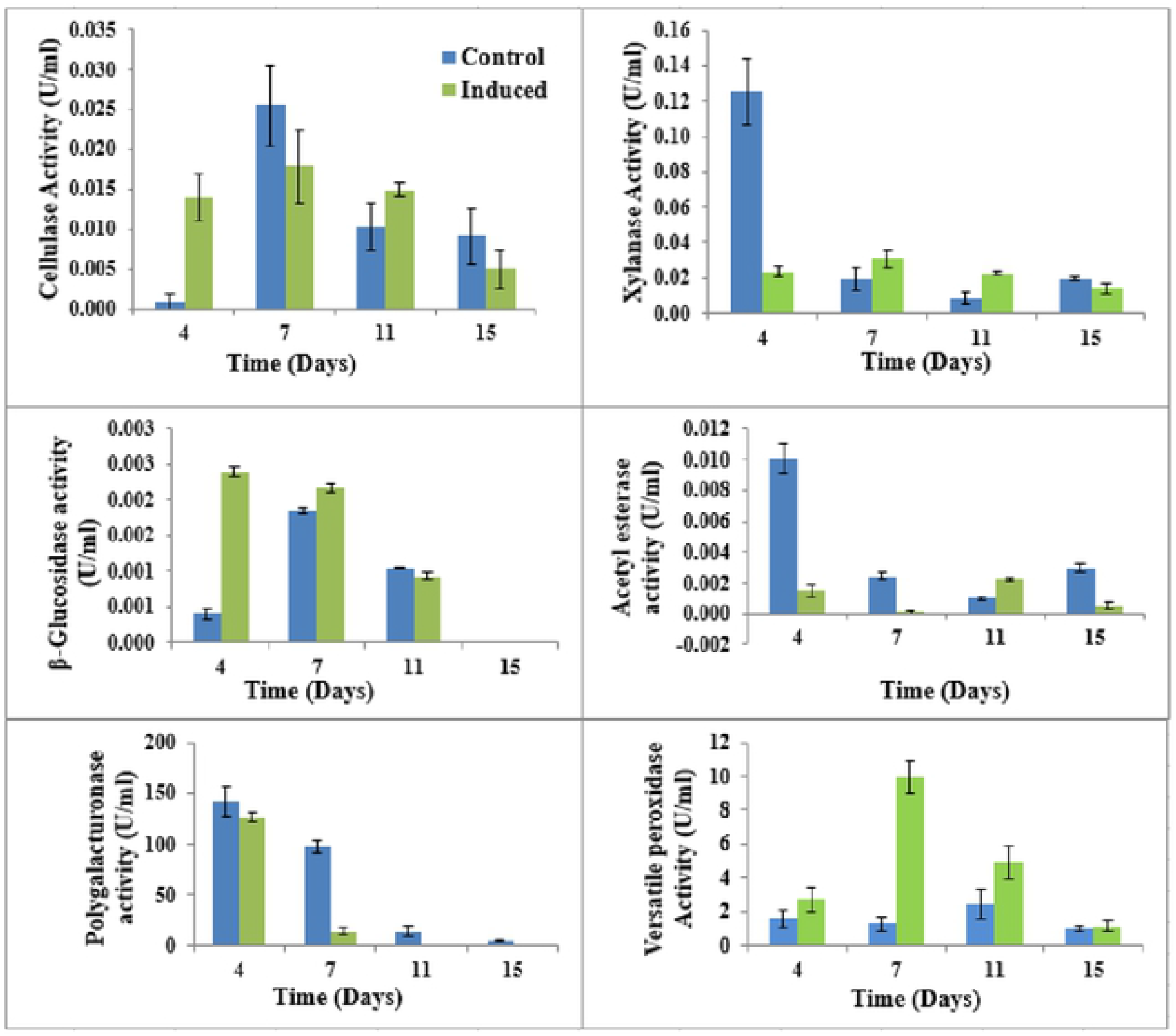
Biochemical analysis of few biomass degrading enzymes in the culture supernatant of *L.squarrosulus*

**Figure 2:**
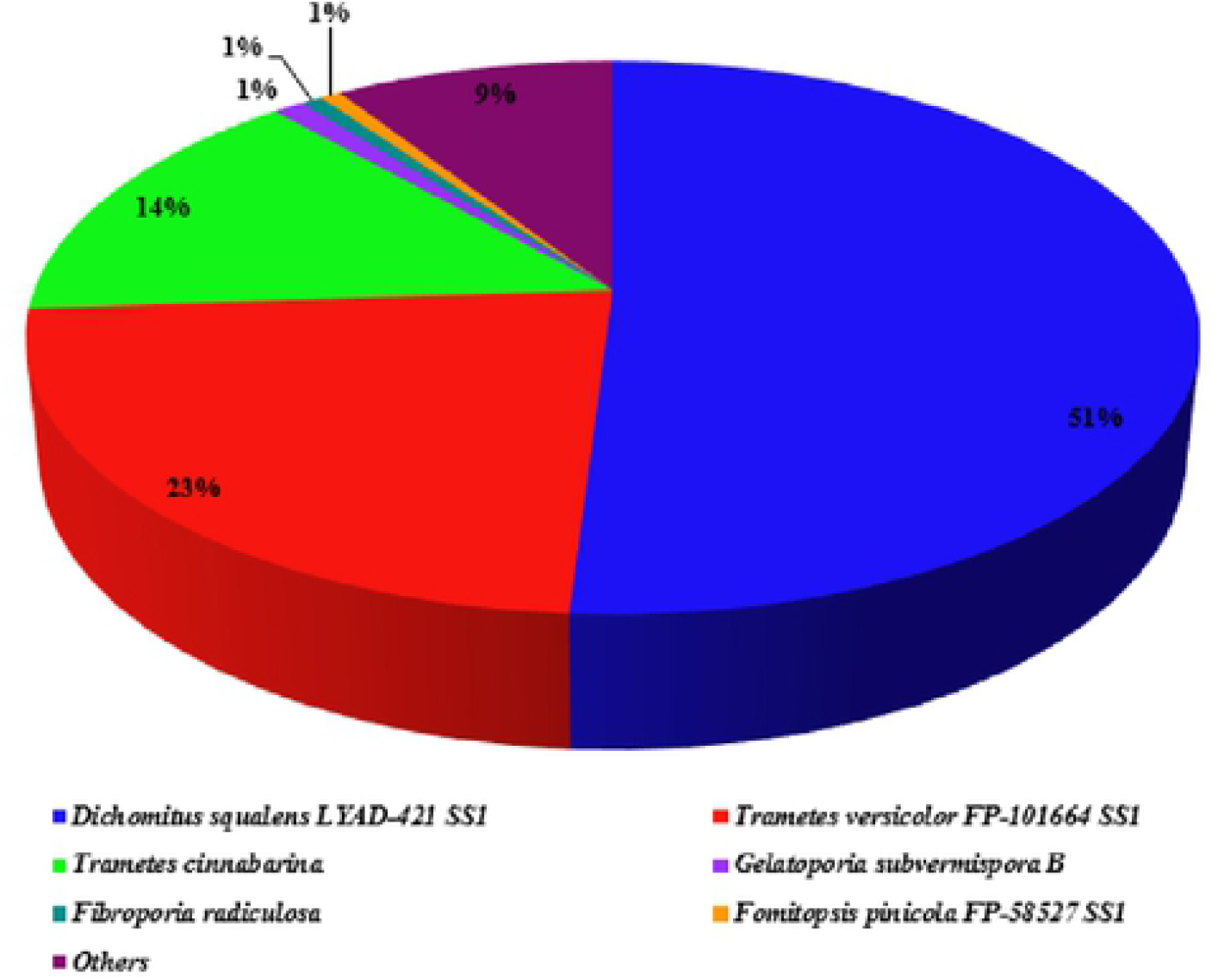
Taxonomic distribution of *L.squarrosulus* nuclcotidescquences at species level.

Gene Ontology terms for the transcripts were annotated through BLAST2GO and Function annotator. 3,365 GO terms were assigned to 10,494 transcripts. The most abundant GO term predicted by Function annotator was GO:0055114, specifying the oxidation-reduction process (biological process) with gene products marking to manganese peroxidase 3 precursor of *Phlebia radiata* (PEM3_PHLRA), laccase 1A of *Trametes pubescens* (AF414808.1, AF491761, AF414807.1), ligninase H2 of *Phanerochaete chrysosporium* (LIG4_PHACH), mannitol dehydrogenase (MTLD_BACP2), NADPH dependent D-xylose reductase (XYL1_CANBO), arabinitol dehydrogenase (ARD1_UROFA), arabinan endo-1,5-alpha-L-arabinosidase A (ABNA_EMENI),pyranose dehydrogenase (PDH3_LEUMG), α-fucosidase A (AFCA_ASPNC). This substantiates the potential of this fungus in biomass degradation through production of diverse hydrolytic enzymes. Subsequently rich GO terms were GO:0005524 (Molecular function:ATP binding), GO:0008152 (Cellular function: Metabolic process) and GO:0016021 (Cellular component: integral membrane components). Deeper classification of each of the GO terms predicted for the contigs were also visualized through Function annotator that presented fifteen levels of classification for each GO category.

Best matching hits of the putative proteins encoded by the transcripts against NCBI non-redundant protein database were available for 16,779 transcripts that expressed similarity to chiefly *Dichomitus* and *Trametes* protein sequences.3,217 probable enzyme products likely to be produced by 12,379 transcripts were determined through PRIAM based on ENZYME database. Though top abundant hits were the proteins involved in genome integrity and regulationlike RNA dependent RNA polymerase, RNA helicase, protein kinases, there were significant representation of biomass degrading enzymes like endo-1,3(4)-beta-glucanase, glucose oxidase, choline oxidase and a range of lignocellulose active enzymes as depicted in Figure 3. This demonstrates that expression of these enzymes by the fungus are not dependent on the presence of lignocellulose substrate in the culture medium.

**Figure 3:**
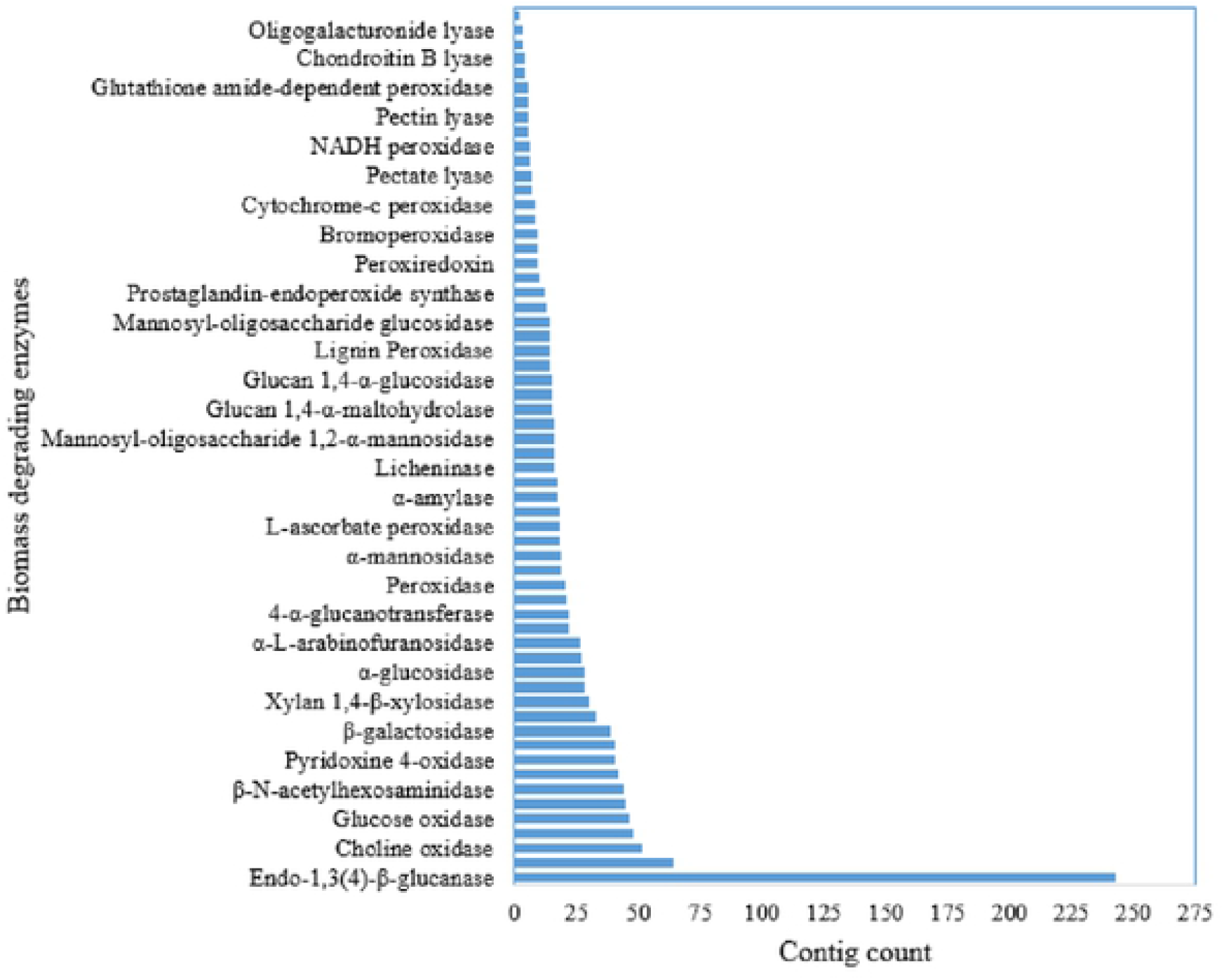
Putative biomass degrading enzymes of *Lentinus squarrosulus*

Putative domain hits illustrated by Function annotator were based on PFAM database. 4952 unique conserved domains were identified against 11,585 transcripts. The most abundant domain hit was Major Facilitator Superfamily of secondary transporters (pfam07690) followed by Tymo_45kd_70kd (pfam03251), a kind of transposable element detected in Basidiomycetes. Similar transposable elements were also reported in *Pleurotus ostreatus*[25].

KEGG orthology designations were obtained for 3,327 transcripts.Transcripts encoding 20 putative cytochrome P450 polypeptides were present in *L.squarrosulus* transcriptome.The probable pathway depiction for cytochrome P450 was metabolism of xenobiotics. In addition, mannosidase, α and β glucosidases, galactosidase, arabinofuranosidase, endoglucanase, pectin esterase, polygalacturonase were identified establishing the biodegradative potential of this fungus.

Besides representation of sequences involved in fungal internal metabolism, translation and transcription, there was considerable expression of two component signal transduction system coupled with transcription factor SKN7. SKN7 transcription factor is an important member of two component phosphorelay system that transfers signal to activate the promoters of genes in response to external stimuli and induces response to oxidative stress such as H_2_O_2_ [26]. In addition, multiple RTA1 domain containing protein sequences were observed, again related to stress response.

There was significant representation of contigs specifying for laccase as revealed through similarity searches to nr database and also by GO annotation. Laccase, a multicopper oxidase catalyzes the oxidation of phenolic compounds using molecular oxygenas electron acceptor as pointed out in the molecular function of the enzyme by GO as hydroquinone: oxygen oxidoreductase activity, hydroquinone being a diphenol compound. Laccases are efficient in degrading the phenolic components of lignin and plays a major role in lignin catabolic process.Six transcriptsof laccase were seen expressed in *L.squarrosulus* with 100% sequencecongruence to laccases of*Trametes cinnabarina*, *Polyporus*, *Lentinus tigrinus*.Ten transcript sequences were found to encode proteins homologous to versatile peroxidase with significant protein similarity to versatile peroxidase protein isoforms of *Pleurotus eryngii* and of *Trametes versicolor*.Protein sequences of putative versatile peroxidases were subjected to multiple alignment through CLUSTALW with experimentally determined protein sequences of ligninolytic peroxidases in Protein Data Bank (PDB). Rooted Phylogenetic tree through UPGMA method of the alignment is presented in Figure 4.

**Figure 4:**
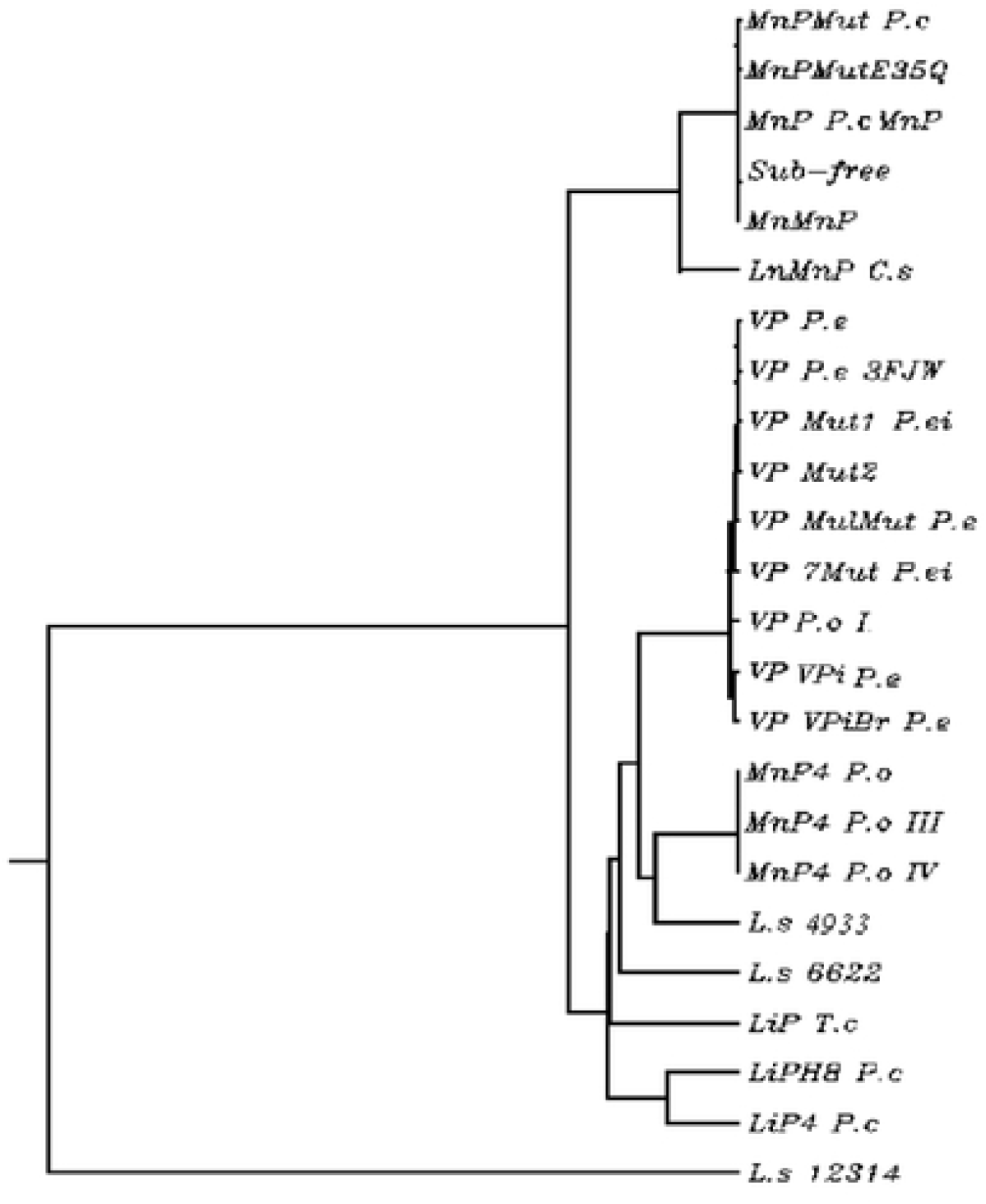
Multiple sequence comparison of Ligninolytic peroxidases with *L.squarrosulus* putative versatile peroxidase sequences [P.c – *Phanerochaete chrysosporium*; C.s – *Ceriporiopsis subvermispora*; P.e – *Pleurotus eryngii,* P.o – *Pleurotus ostreatus*; T.c – *Trametes cervina*; L.s – *Lentinus squarrosulus*]

Two isoforms of manganese peroxidase were identified based on the conserved domains and sequence similarity. However, there were no lignin peroxidase transcripts observed with this species as confirmed through the biochemical analysis.

Peroxidases are more prominent in the lignin catabolic process for their relatively high redox potential and hence requisite for complete mineralization of lignin. Efficient function of ligninolytic machinery is indispensable without hydrogen peroxide, electron acceptor for peroxidases. White-rot fungi produce H_2_O_2_ needed for lignin oxidation through diverse enzymes like glyoxal oxidase, aryl-alcohol oxidase, aryl alcohol dehydrogenase and GMC oxidoreductases [27]. Glyoxal oxidase of this fungus exhibited 66% similarity in protein sequence to that of *Phanerochaete chrysosporium*. Multiple transcript copies of these enzymes were seen expressed by the fungus.A considerable depiction of H_2_O_2_ producing oxidases were made by Glucose-Methanol-Choline (GMC) superfamily encoding transcripts which supplies the peroxide requisite for ligninolytic peroxidases.

Enhancement of oxido-reduction process in the culture of this fungus was further validated by ubiquitous presence of Cytochrome P450 monooxygenases and oxidoreductases in the transcriptome [28, 29]. Cytochrome P450 enzymes are involved in the catalysis reaction of aromatics metabolism through diverse biochemical reactions. The enzymatic reaction of cleavage of β-O-4 linkages of lignin are enhanced by co-oxidants like thiols, NAD^+^, glutathione etc. Glutathione reductase enables regeneration of reduced glutathione and is seen expressed in cells exposed to oxidative stress [30]. Evidently Glutathione reductase was seen expressed in *L.squarrosulus* with considerable protein similarity to glutathione reductase of *Dichomitus squalens*. In addition, sequences of flavin containing monooxygenases (FMO) and dioxygenases were revealed in the analysis of the transcriptome. FMOs are proteins involve in degradation in multitude of aromatics and have been reported in a number of fungal species while dioxygenases are reportedly involved in ring cleavage of aromatics oxidation [31]. The reactive peroxides produced create a highly oxidative environment for enzymatic action. Fungi secrete svf1 protein in response to this oxidative stress for its survival as evident through the sequences of oxidative stress survival svf1-like protein seen expressed by *L.squarrosulus* [32].

Plant polysaccharides are composed of cellulose, hemicellulose, pectin and lignin which contribute to the bulk of the biomass. Besides the expression of lignin degrading enzymes in response to induction by aromatic compounds, polysaccharide degrading enzymes also existed in the transcriptome with regulation different from that of control mediumindicating a synergy in regulation of the former and the latter. Endoglucanases, cellobiohydrolases and β-glucosidases enzymes are important components of the cellulolytic machinery [33] and *L.squarrosulus* demonstrated the presence of mRNAs of these genes. Pectin lyase like proteins belong to glycosyl hydrolase family 28 that act by inverting mechanism on α1,4 glycosydic linkage of the polygalacturonates [34]. Sequences with pectin lyase like protein activity belonging to pectin esterase, polygalacturonase in the transcriptome of this species state its role in degradation of pectin.Besides pectin lyases, transcripts of polysaccharide lyase classes of protein were also seen expressed. In addition, there was strong representation of substrate transporters, glycoside hydrolases, glycoside transferases, carbohydrate esterases and acetyl xylan esterases encompassing the major plant polysaccharide degrading enzymes.

### CAZyme annotation

The fungus undertaken in the current study was an extensive producer of multitude of plant polysaccharide degrading enzymes spanning across the families of carbohydrate active enzymes.Fungal systems are reportedly affluent in polysaccharide degrading enzymes and the enzymes acting on these polysaccharides in general are designated to 156 families of glycoside hydrolases, 106 families of glycosyl transferases, 16 families of carbohydrate esterases and 29 families of polysaccharide lyases [35]. The transcriptome was found enriched with transcripts of esterases of family carbohydrate esterases 10. Carbohydrate esterases deacetylate the conjugates of glucans and are binding components of polysaccharide degrading machinery. Glycoside hydrolases of family 16 showed significant depiction in the transcriptome of *L.squarrosulus*. Also observed were the transcript sequences of six and seven hairpin glycosidases catalyzing O-glycosyl bonds. This emphasizes the importance of this fungus in catabolism of carbohydrates majorly cellulose, hemicellulose and pectic polysaccharides. In this bracket, endo glucanases, β 1, 3-1, 4 glucanases, xyloglucanases, xyloglucan: xyloglucosyl transferases worth remarkable mention.

Chitinases of glycoside hydrolases family 18 was subsequently predominant that was supposed to be involved in cell wall remodeling and maintenance of the fungus. More than ten transcripts with GH3 and GH79 modules were identified in the transcriptome of *L.squarrosulus*. Similarly, ligninolytic enzymes that act in synchronization with the glycoside hydrolases were also preponderant in the transcriptome as revealed by CAZyme annotation belonging to AA2 family. The other affluent CAZymes reported were GMC oxidoreductase (AA3), lytic polysaccharide monooxygenases cleaving cellulose chains (AA9) and lytic polysaccharide monooxygenases cleaving xylans (AA14).Overall distribution of CAZymes in the transcriptome based on dbCAN is illustrated in Figure 5-7.

**Figure 5:**
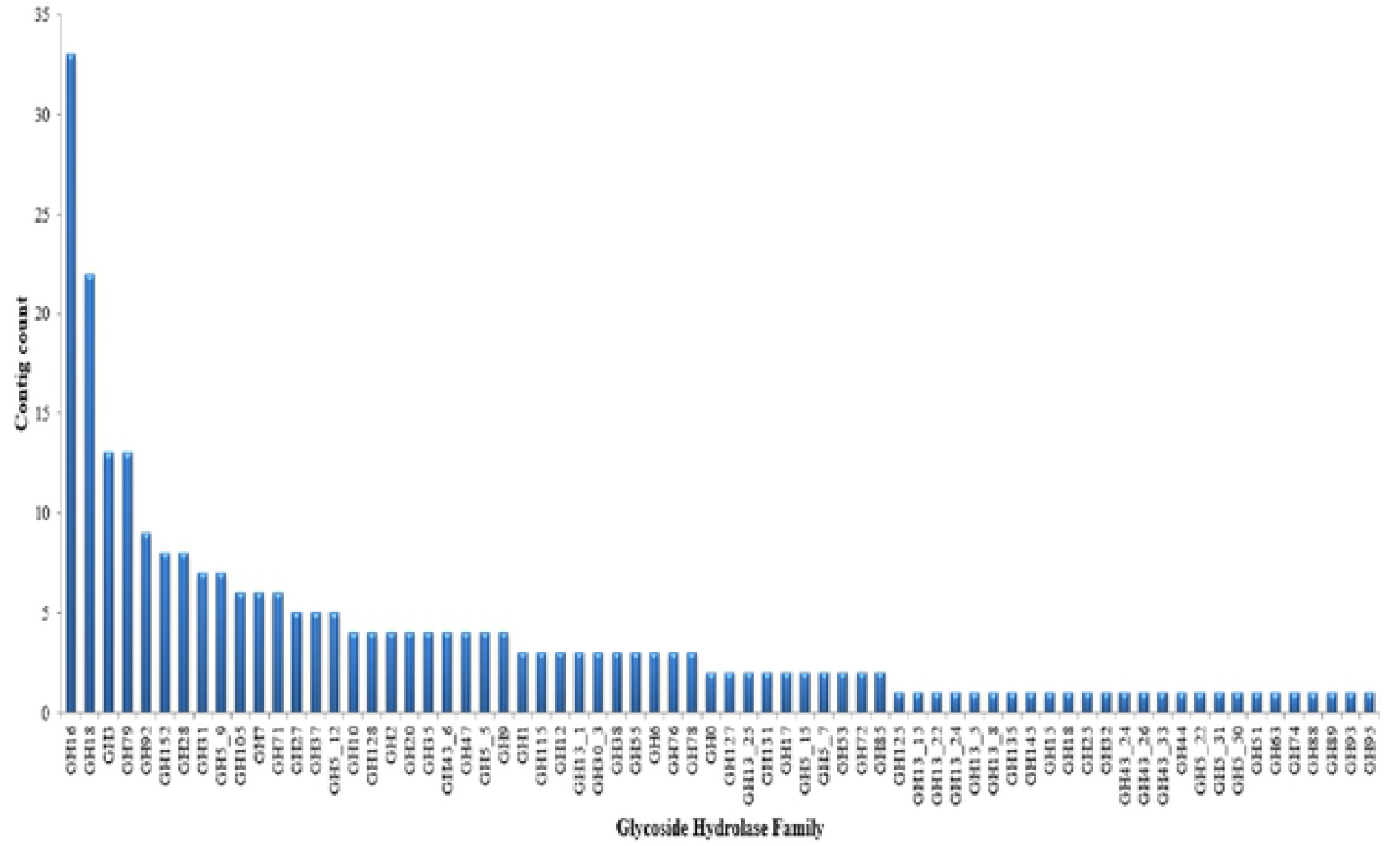
Distribution of Glycoside hydrolases (GH) in the transcriptome predicted based on CAZyme database.

**Figure 6:**
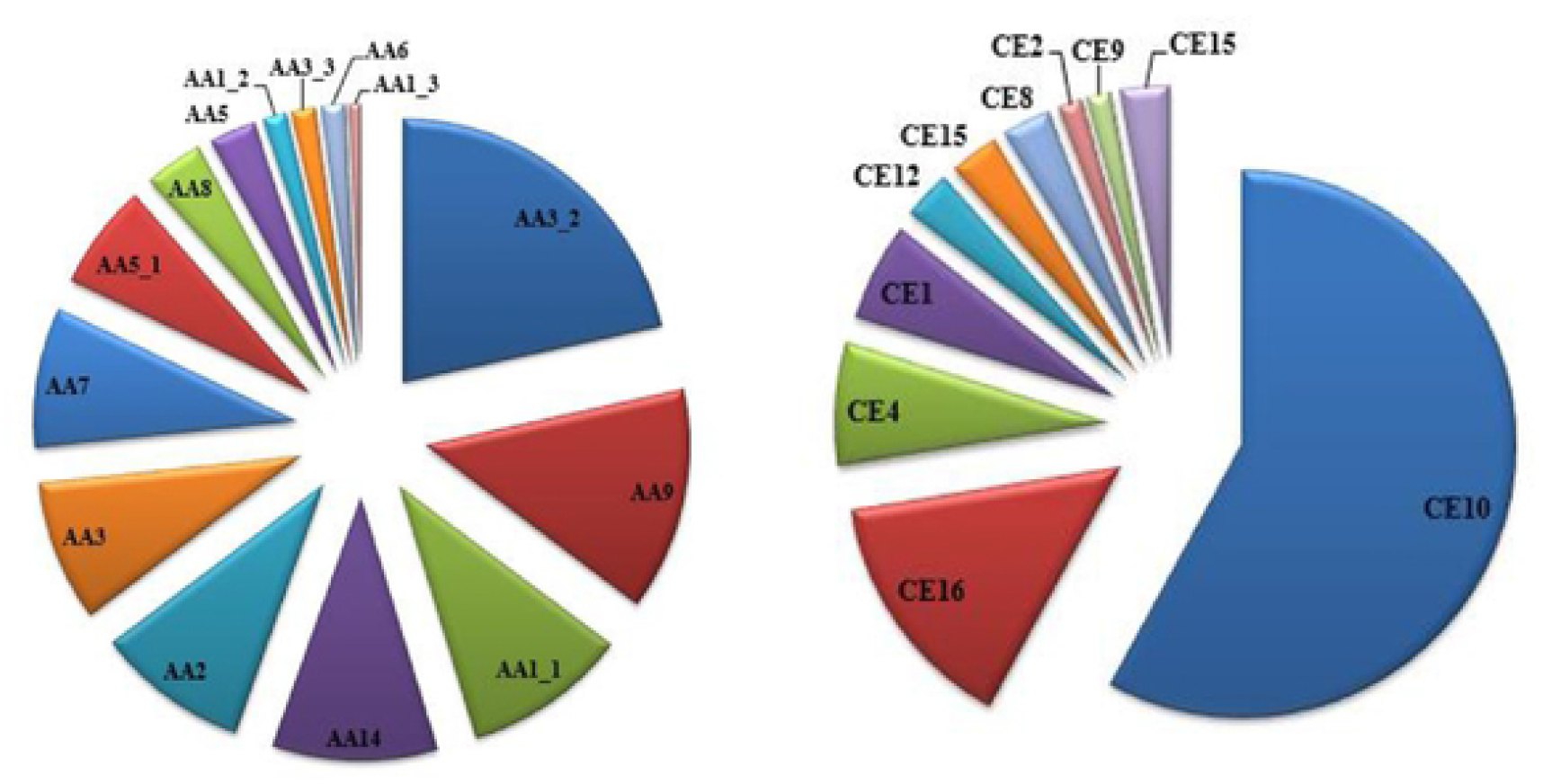
Distribution of Auxiliary activities (AA) enzymes and carbohydrate esterases (CE) in the transcriptome predicted based on CAZyme database.

**Figure 7:**
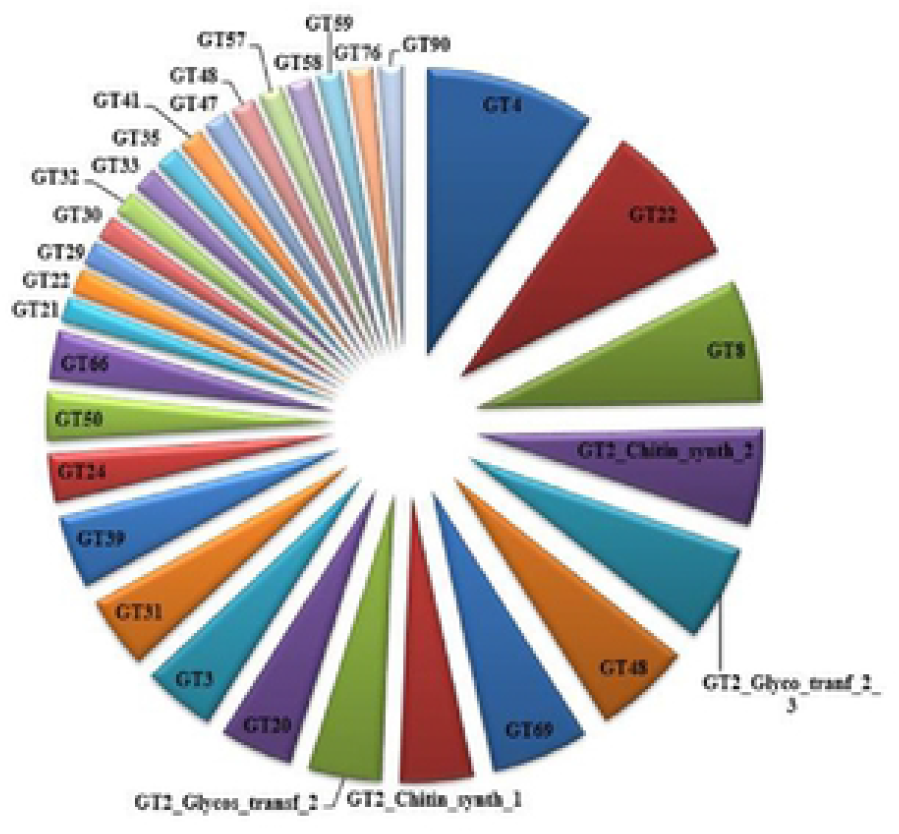
Distribution of glycosyl transferases (GT) in the transcriptomc predicted based on CAZyme database.

CUPP assigned 508 transcripts to 6 families of polysaccharide lyases, 51 families of glycoside hydrolases, 23 families of glycoside transferases, 7 families of carbohydrate esterases and 10 families of auxiliary activities family.Of the enzymes characterized through CUPP based on the peptide signatures, chitinase (3.2.1.14)and chitin synthase (2.4.1.16)was predominant followed by laccase (1.10.3.2). Ten transcript sequences were assigned to laccase family and seven transcripts encoded peptide signatures typical of versatile peroxidase (1.11.1.16). The top hits also included glyoxal oxidase (1.2.3.15) and glucan endo-1,3-beta-D-glucosidase (3.2.1.39). Prediction of CAZy families on peptide signatures revealed preponderance of glycosidehydrolases followed by auxiliary activities 3 family. Among the glycoside hydrolases, GH16 and GH18 were enriched in the transcript sequences of *L.squarrosulus.* GH16 comprises of members that are active on β-1,4 and 1,3 glycosidic bonds and GH18 members chitinase and endo-β-N-acetylglucosaminidase aid in maintenance of fungal cell wall whereasGH5 activities include cellulases, endomannanases and xyloglucanases.

Auxiliary activities 2 family encoding lignin modifying peroxidases manganese peroxidase and versatile peroxidase were also abundant subsequent to glycoside hydrolases and auxiliary activities family 3.

## MycoCLAP ANALYSIS

Homology search results against the mycoCLAP nucleic acid sequences database of approximately 833 sequences functioning on lignocellulose degradation are depicted in Table 2. Ranking of search results were based on closest biochemically characterized homologue.

**Table 1:** Summary of *L.squarrosulus* transcriptome assembly

|  |  |
| --- | --- |
| Number of contigs | 25, 244 |
| Total size of contigs (bp) | 2,31,81,074 |
| Average Length (bp) | 918 |
| Length SD (bp) | 856 |
| GC content (%) | 57.93 |
| N50 contig length (bp) | 1,340 |
| N50 contig count | 5,091 |

**Table 2:**
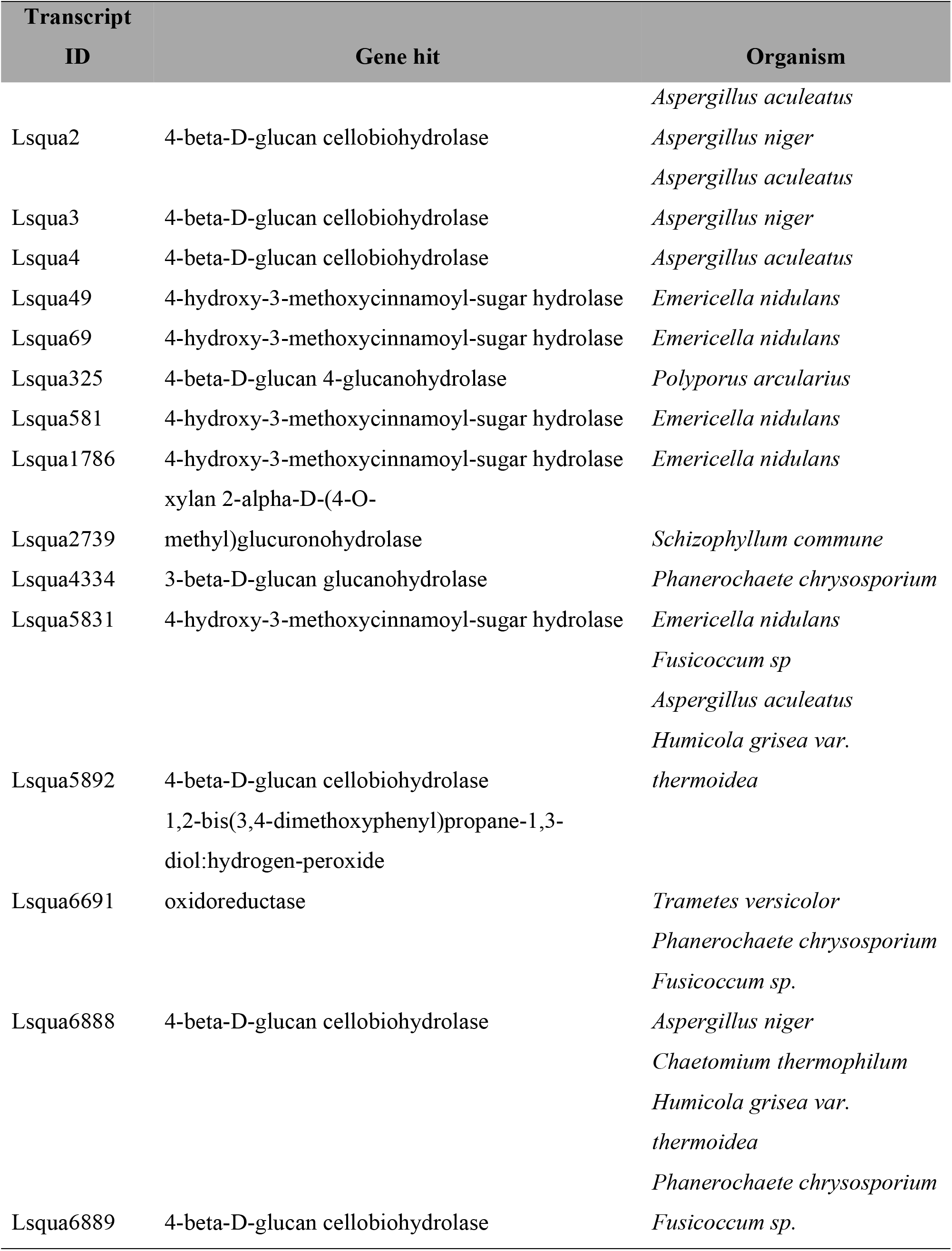
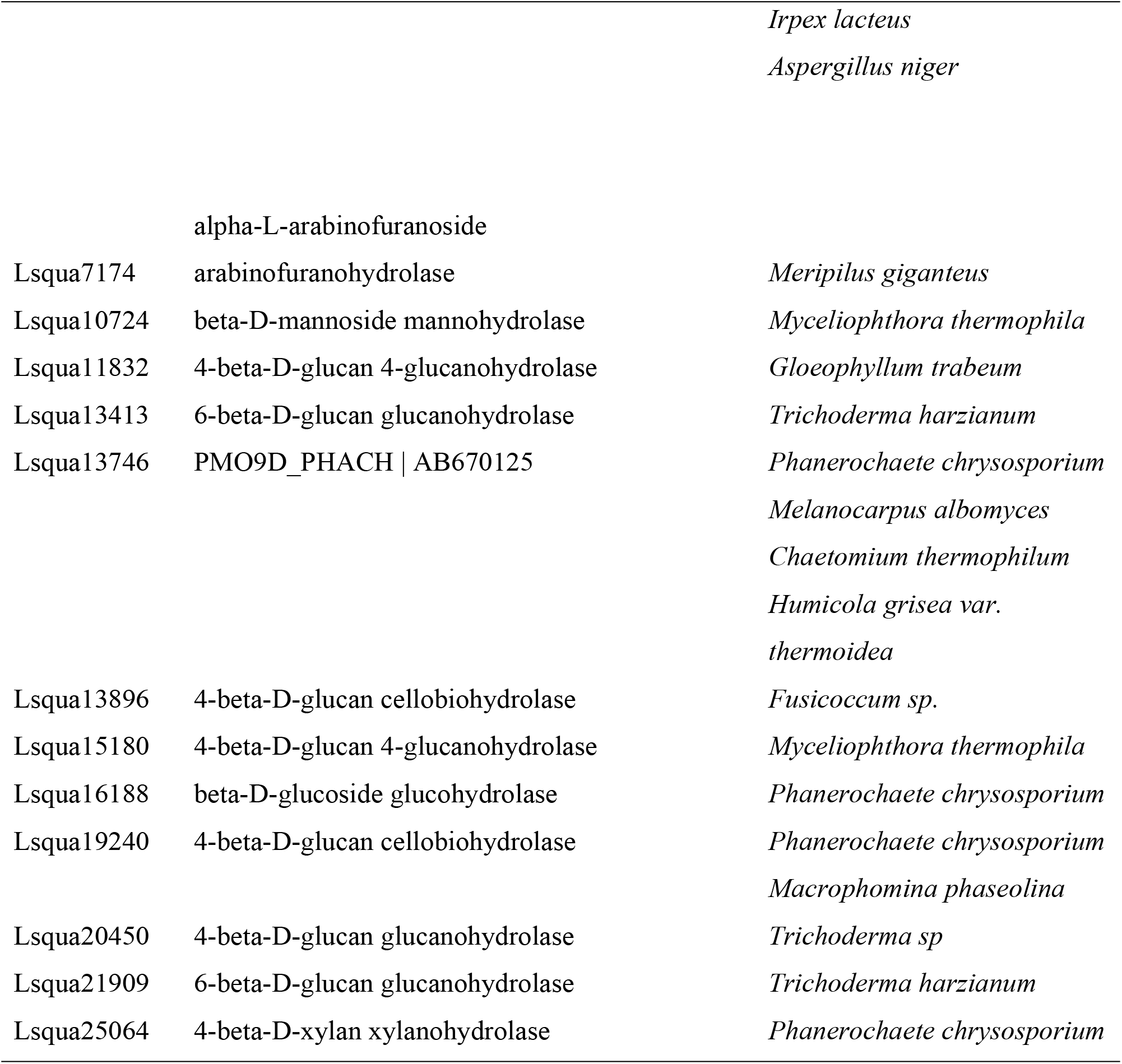
Nucleotide hits against mycoCLAP database.

### Gene prediction

The predicted proteins of six transcripts corresponded to laccases, the multicopper oxidases. The protein sequences predicted by AUGUSTUS and FGENESH were similar except for slight length variations for a few transcripts. Laccase expression was visualized in the medium devoid of external supplementation of copper in contrast to cultures requiring Cu^2+^ addition for laccase induction [36, 37]. Though two of the sequences were only partial, three sequences encoded proteins with length > 399 aa. The predicted proteins exhibited more than 90% similarity to laccases of *Lentinus tigrinus* and *Polyporus brumalis*. The sequences exhibited conserved L1-L4 signatures typical of laccases[38] involved in copper ion binding. Ten transcripts were identified to encode product of versatile peroxidase as assessed through aminoacid sequence alignment. The sequences exhibited significant homology with manganese dependent and repressed peroxidase isoforms of *Lentinus tigrinus* and *Trametes versicolor*.Some sequences though partial presented heme, metal ion binding domains.A complete versatile peroxidase coding sequence was also predicted with 92% protein identity to manganese peroxidase 1 isoform of *Lentinus tigrinus*. The predicted proteins did include fatty acid dehydrogenases involved in peroxidation of lipids which generates lipid radicals as ligninolytic oxidants for non-phenolic lignin degradation by manganese peroxidases.Protein sequence comparison of predicted ligninolytic enzymes is illustrated in Table 3.

**Table 3:** Putative ligninolytic enzymes of *L.squarrosulus*

| Transcript ID | Putative Function | BLAST best hit | Sequence length | e-value | Amino acid identity |
| --- | --- | --- | --- | --- | --- |
| Lsqua4933 | Versatile Peroxidase | manganese peroxidase 1 [*Lentinus tigrinus*] | 369aa | 0 | 92% |
| Lsqua6622 | Versatile Peroxidase | manganese-repressed peroxidase [*Trametes versicolor*] | 280aa | 4e<sup>-139</sup> | 72% |
| Lsqua18051 | Versatile Peroxidase | manganese peroxidase isozyme precursor [*Polyporus brumalis*] | 179aa | 8e<sup>-117</sup> | 94% |
| Lsqua10154 | Versatile Peroxidase | Mn peroxidase MNP6<br>[*Lentinus tigrinus*] | 168aa | 1e<sup>-109</sup> | 96% |
| Lsqua13135 | Versatile Peroxidase | manganese peroxidase<br>isozyme precursor<br>[*Lentinus tigrinus*] | 72aa | 7e<sup>-35</sup> | 86% |
| Lsqua15894 | Versatile Peroxidase | manganese peroxidase<br>isozyme precursor<br>[*Lentinus tigrinus*] | 65aa | 1e<sup>-31</sup> | 92% |
| Lsqua15340 | Versatile Peroxidase | manganese peroxidase<br>isozyme precursor<br>[*Lentinus tigrinus*] | 135aa | 2e<sup>-84</sup> | 93% |
| Lsqua10666 | Versatile Peroxidase | Mn peroxidase MNP6<br>[*Lentinus tigrinus*] | 76aa | 3e<sup>-38</sup> | 87% |
| Lsqua14505 | Versatile Peroxidase | Mn peroxidase MNP6<br>[*Lentinus tigrinus*] | 76aa | 3e<sup>-38</sup> | 87% |
| Lsqua12314 | Versatile Peroxidase | heme peroxidase [*Lentinus tigrinus*] | 406aa | 0 | 80% |
| Lsqua20933 | Laccase | laccase [*Lentinus tigrinus*] | 68aa | 9e<sup>-24</sup> | 72% |
| Lsqua6164 | Laccase | laccase 1 [*Polyporus*<br>*brumalis*] | 516aa | 0 | 88% |
| Lsqua18153 | Laccase | laccase [*Polyporus*<br>*brumalis*] | 474aa | 0 | 92% |
| Lsqua18367 | Laccase | laccase LCC3-3<br>[*Polyporus ciliatus*] | 399aa | 0 | 93% |
| Lsqua21132 | Laccase | Cu-oxidase-domain-<br>containing protein<br>[*Lentinus tigrinus*] | 133aa | 3e<sup>-84</sup> | 96% |
| Lsqua21539 | Laccase | laccase [*Lentinus tigrinus*] | 244aa | 1e<sup>-149</sup> | 91% |

## Discussion

Basidiomycetes, particularly white-rot fungi have been investigated exhaustively for their selective delignification ability and also for efficiency of lignin degradation. These fungi degrade plant biomass through a repertoire of biomass degrading enzymes. On these lines there had been tremendous effort to deepen the understanding of the biomass metabolic process especially the lignin catabolic process as degradation of lignin will facilitate competent utilization of the underlying polysaccharides. Ligninolytic enzymes are reported to be secreted in response to stress such as nutrient limitation, presence of recalcitrant aromatic compounds and also triggered by presence of metal ions such as Cu^2+^ and Mn^2+^ in case of laccase and manganese peroxidase respectively [4, 36, 38, 39]. White-rot fungal species vary widely in their ability to secrete these biomass degrading enzymes and the enzyme production itself is largely influenced by growth conditions and presence of substrates [40]. With the genomic data of white-rot fungal species accumulating, characterization of the biomass degrading enzymes for functional exploitation has increased. Transcriptomic studies have also revealed the expression of an array of catabolic enzymes differentially based on the lignocellulosic substrate [4, 5, 6, 9, 41]. Accordingly in our study, we studied the transcriptome of *L.squarrosulus,* a white-rot belonging to Polyporaceae family grown in simple culture medium with azo dye Reactive Black 5 (RB5) added for ligninolytic enzyme induction. Our study was focused to study the expression of degradative enzymes in simple medium and not intended for differential expression analysis. However, the transcriptomic profile of this fungus correlated well with the transcriptomic data of other white-rot fungi available [6, 8, 42]. There was significant expression of CAZymes especially ligninolytic enzymes in the induced medium in support of the fact that ligninolytic enzymes are mainly expressed in nutrient limited conditions rather than presence of lignocellulosic substrates.

Multiomics studies on *Phanerochaete chrysosporium* lignocellulolytic networks also substantiated that nutrient limitation were major drivers of ligninolytic and cellulolytic gene expression rather than the presence of lignocellulosic substrates [4]. Cellulase, xylanase, polygalacturonase, β-glucosidase, laccase and versatile peroxidase are shown to be produced in considerable amounts despite the lack of lignocellulosic substrate. A noteworthy observation is increase in amount of ligninolytic enzymes secreted in comparison to control medium. Putative cellulases (EC 3.2.1.4) and xylanases (EC 3.2.1.8) were identified in the transcriptome of our fungus in spite of presence of glucose in the medium which is presumed to cause catabolite repression.However, glucose concentration in the medium during the period of RNA isolation was comparatively low that might have plausibly caused expression of cellulose degrading enzymes. In addition, transcripts encoding putative hemicellulose degrading enzymes of arabinanase, xylosidase, mannanase were also seen expressed in the transcriptome of *L.squarrosulus*. These results are in contrast to the transcriptomic analysis of *Pycnoporus sanguineus* also grown in the absence of lignocellulosic substrate wherein cellulases were not present [42]. However, in our present study, cellulases and hemicellulases were perceived to be expressed even in the absence of lignocellulosic substrates.In addition,lytic polysaccharide monooxygenases (LPMO), pectinases and a range of other polysaccharide degrading enzymes were detected elucidating the superior biodegradative ability of the fungus. Similar oxidative enzymes acting on polysaccharides were seen expressed in *Ceriporiopsissubvermispora* and *Phanerochaete chrysosporium*cultures containing lignocellulosic substrates [43, 44]. Ligninolysis is substantially connected to free radical and peroxide production. *L.squarrosulus* transcriptome revealed diverse peroxide generating enzymes like cellobiose dehydrogenases, aryl alcohol oxidases and other members of glucose-methanol-choline oxidoreductases. In a nutshell, the fungus undertaken in our study, *L.squarrosulus* is evidently competent to efficiently degrade lignocelluloses through its extensive machinery of hydrolytic and oxidative enzymes targeting multiple components of the substrate. Additionally, these enzymes working on lignocellulose bioconversion also react on diverse aromatic compounds like dyes, pesticides, endocrine disruptors and other environmental contaminants which convey their importance. Though multiple white-rot Basidiomycetes were studied for lignin degradation, each species is unique in its ability to oxidize the aromatic macromolecule.

### Conclusion

Though there are multiple studies on biochemical characterization of ligno-cellulolytic enzymes of *L.squarrosulus,* research on expression pattern of the lignocellulose degrading enzymes has not been attempted. This study on the transcriptome analysis of *L.squarrosulus* revealed significant facts on this front and will definitely enhance the knowledge about the biodegradative ability of this fungus potentially paving the way for efficient biotechnological applications utilizing its potency in biomass degradation and its future functional exploitation in biomass conversion applications.

## Acknowledgements

The financial assistance by Department of Science and Technology (DST), Ministry of Science and Technology, Govt. of India, under the WOSA scheme (SR/WOS-A/LS-32/2016) is gratefully acknowledged by the first author. The authors thank the Director, ICAR - National Institute of Animal Nutrition and Physiology, Bangalore (Karnataka) India, for providing the necessary facilities to carry out the research work.

## References

1. Valverde ME, Hernández-Pérez T, and Paredes-López O (2015) Edible Mushrooms: Improving Human Health and Promoting Quality Life. Int J of Microbiol:14.

2. Kent KT (1987) Enzymatic ‘combustion’, the microbial degradation of Lignin. Ann. Rev. Microbiol 41: 465–505.

3. Sridhar M and Ravichandran A (2017) Insights into the mechanism of lignocellulose degradation by versatile peroxidases. Curr sci 113:35–42.

4. Wymelenberg AV, Gaskell J, Mozuch M, Kersten P, Sabat G, Martinez D, Cullen D (2009) Transcriptome and secretome analyses of *Phanerochaete chrysosporium* reveal complex patterns of gene expression. Appl Environ Microbiol 5(12):4058–68.

5. Chen Y, Cao Q, Tao X, Shao H, Zhang K, Zhang YZ and Tan X (2016) Analysis of *de novo* sequencing and transcriptome assembly and lignocellulolytic enzymes gene expression of *Coriolopsis gallica* HTC. Biosci Biotechnol Biochem 81(3):460–468.

6. Zhang L, Wang Z, Wang Y, Huang B (2017) Transcriptomic profile of lignocellulose degradation from *Trametes versicolor* on poplar wood. Bioresources 12(2):2507–2527.

7. Makelä MR, Sietiö O-M, de Vries RP, Timonen S, Hildén K (2014) Oxalate-Metabolising Genes of the White-Rot Fungus *Dichomitus squalens* Are Differentially Induced on Wood and at High Proton Concentration. PLoS ONE 9(2): e87959.

8. Korripally P, Hunt CG, Houtman CJ, Jones DC, Kitin PJ, Cullen D and Hammel KE (2015) Regulation of gene expression during the onset of ligninolytic oxidation by *Phanerochaete chrysosporium* on spruce wood. Appl Environ Microbiol 81:7802–7812.

9. Qin X, Su X, Luo H, Ma R, Yao B and Ma F (2018) Deciphering lignocellulose deconstruction by the white rot fungus *Irpex lacteus* based on genomic and transcriptomic analyses. Biotechnol Biofuels 11:58.

10. Shewale JG and Sadana JC (1978) Cellulase and β-glucosidase production by a basidiomycete species. Can J Microbiol 24(10):1204–16.

11. Li Q, Coffman AM and Ju LK (2015) Development of reproducible assays for polygalacturonase and pectinase. Enzyme Microb Technol 72:42–8.

12. Kataoka N and Tokiwa Y (1998) Isolation and characterization of an active mannanase producing anaerobic bacterium, *Clostridium tertium* KT-5A, from lotus soil. J Appl Microbiol 84(3):357–67.

13. Miller GL (1959) Use of Dinitrosalicylic Acid Reagent for Determination of Reducing Sugar. Anal Chem 31(3): 426–428.

14. Biely P, Puls J and Schneidera H (1985) Acetyl xylan esterases in fungal cellulolytic systems. FEBS Lett 186(1):80–84.

15. Wariishi H, Valli K and Gold MH (1992) Manganese(II) Oxidation by Manganese Peroxidase from the Basidiomycete *Phanerochaete chrysosporium.* Kinetic mechanism and role of Chelators. J Biol Chem 267(33):23688–95.

16. Conesa A, Götz S, García-Gómez JM, Terol J, Talón M and Robles M (2005) Blast2GO: a universal tool for annotation, visualization and analysis in functional genomics research. Bioinformatics 21(18):3674–6.

17. Chen TW, Gan RC, Fang YK, Chien KY, Liao WC, Chen CC, Wu TH, Chang IY, Yang C, Huang PJ, Yeh YM, Chiu CH, Huang TW and Tang P (2017) FunctionAnnotator, a versatile and efficient web tool for non-model organism annotation. Sci Rep 7(1):10430.

18. Moriya Y, Itoh M, Okuda S, Yoshizawa AC and Kanehisa M (2007) KAAS: an automatic genome annotation and pathway reconstruction server. Nucleic Acids Res 35: W182–5.

19. Zhang H, Yohe T, Huang L, Entwistle S, Wu P, Yang Z, Busk PK, Xu Y and Yin Y (2018) dbCAN2: a meta server for automated carbohydrate-active enzyme annotation. Nucleic Acids Res 46(W1):W95–W101.

20. Barrett K, Lange L (2019) Peptide-based functional annotation of carbohydrate-active enzymes by conserved unique peptide patterns (CUPP). Biotechnol Biofuels 12, 102.

21. Strasser K, McDonnell E, Nyaga C, Wu M, Wu S, Almeida H, Meurs MJ, Kosseim L, Powlowski J, Butler G and Tsang A (2015) mycoCLAP, the database for characterized lignocellulose-active proteins of fungal origin:resource and text mining curation support. Database (Oxford): bav008.

22. Stanke M and Morgenstern B (2005) AUGUSTUS: a web server for gene prediction in eukaryotes that allows user-defined constraints. Nucleic Acids Res 33:W465–7.

23. Solovyev V, Kosarev P, Seledsov I and Vorobyev D (2006) Automatic annotation of eukaryotic genes, pseudogenes and promoters. Genome Biol: 10.1-10.12.

24. Alfredo J, Otto M, Dimitrios F, Beatriz O, Elisabet S, Daniel L, Karen N, Tuomo N, Karl-Henrik L, Leif R, David S. Hibbett (2017) A revised family-level classification of the Polyporales (Basidiomycota). Fungal Biol 121(9) 798–824.

25. Andres RC (2017) Transposable elements in basidiomycete fungi dynamics and impact on genome architecture and transcriptional profiles, PhD thesis, public university of Navaraa.

26. Lee J, Godon C, Lagniel G, Spector D, Garin J, Labarre J and Toledano MB (1999) Yap1 and Skn7 Control Two Specialized Oxidative Stress Response Regulons in Yeast. J Biol Chem 274: 16040.

27. Ferreira P, Carro J, Serrano A and Martínez AT (2015) A survey of genes encoding H2O2-producing GMC oxidoreductases in 10 Polyporales genomes. Mycologia 107(6):1105–19.

28. Crešnar B and Petrič S (2011) Cytochrome P450 enzymes in the fungal kingdom. Biochim Biophys Acta 1814(1):29–35.

29. Floudas D1, Binder M, Riley R, Barry K, et al. (2012) The Paleozoic origin of enzymatic lignin decomposition reconstructed from 31 fungal genomes. Science 336:1715–1719.

30. Pollegioni L, Tonin F and Rosini, E. (2015) Lignin-degrading enzymes. FEBS J, 282: 1190–1213.

31. Eswaramoorthy S, Bonanno JB, Burley SK and Swaminathan S (2006) Mechanism of action of a flavin containing monooxygenase. Proc Natl Acad Sci U S A 103(26):9832–7.

32. Brace JL, Vanderweele DJ, Rudin CM (2005) Svf1 inhibits reactive oxygen species generation and promotes survival under conditions of oxidative stress in Saccharomyces cerevisiae. Yeast 22(8):641–52.

33. Baldrian P and Valásková V (2008) Degradation of cellulose by basidiomycetous fungi. FEMS Microbiol Rev 32(3): 501–521.

34. van den Brink J, de Vries RP (2011) Fungal enzyme sets for plant polysaccharide degradation. Appl Microbiol Biotechnol 91(6):1477–92.

35. Lombard V, Golaconda Ramulu H, Drula E, Coutinho PM, Henrissat B (2014) The Carbohydrate-active enzymes database (CAZy) in 2013. Nucleic Acids Res 42:D490–D495.

36. Palmieri G, Giardina P, Bianco C, Fontanella B, Sannia G. (2000) Copper induction of laccase isoenzymes in the ligninolytic fungus *Pleurotus ostreatus*. Appl Environ Microbiol 66(3):920–4.

37. Kumar SV, Phale PS, Durani S and Wangikar PP (2003) Combined sequence and structure analysis of the fungal laccase family. Biotechnol Bioeng 83: 386–394.

38. Fernandez-Fueyo E, Ruiz-Dueñas FJ, Ferreira P, Floudas D, Hibbett DS, et al. (2012) A Comparative genomics of *Ceriporiopsis subvermispora* and *Phanerochaete chrysosporium* provide insight into selective ligninolysis. Proc Natl Acad Sci USA 109: 5458–5463.

39. Jeffries TW, Choi S, Kirk TK. (1981) Nutritional regulation of lignin degradation by *Phanerochaete chrysosporium*. Appl Environ Microbiol 42: 290–296.

40. Janusz G, Kucharzyk KH, Pawlik A, Staszczak M and Paszczynski AJ (2013) Fungal laccase, manganese peroxidase and lignin peroxidase: Gene expression and regulation. Enzyme Microb Technol 52(1):1–12.

41. Yakovlev IA, Hietala AM, Courty PE, Lundell T, Solheim H and Fossdal CG (2013) Genes associated with lignin degradation in the polyphagous white-rot pathogen *Heterobasidion irregulare* show substrate-specific regulation. Fungal Genet Biol 56:17–24.

42. Rohr CO, Levin LN, Mentaberry AN, Wirth SA (2013) A First Insight into Pycnoporus sanguineus BAFC 2126 Transcriptome. PLoS One 8(12): e81033.

43. Hori C, Gaskell J, Igarashi K, Kersten P, Mozuch M, Samejima M and Cullen D (2014) Temporal Alterations in the Secretome of the Selective Ligninolytic Fungus *Ceriporiopsis subvermispora* during Growth on Aspen Wood Reveal This Organism’s Strategy for Degrading Lignocellulose. Appl Environ Microbiol 80(7):2062–70.

44. Kameshwar AK and Qin W (2017) Metadata Analysis of *Phanerochaete chrysosporium* Gene Expression Data Identified Common CAZymes Encoding Gene Expression Profiles Involved in Cellulose and Hemicellulose Degradation. Int J Biol Sci 13(1): 85–99.

